# Chemokine profiling of melanoma-macrophage crosstalk identifies CCL8 and CCL15 as prognostic factors in cutaneous melanoma

**DOI:** 10.1101/2023.10.04.560856

**Authors:** Celia Barrio-Alonso, Alicia Nieto-Valle, Elena García-Martínez, Alba Gutiérrez-Seijo, Verónica Parra-Blanco, Iván Márquez-Rodas, José Antonio Avilés-Izquierdo, Paloma Sánchez-Mateos, Rafael Samaniego

## Abstract

During cancer evolution, tumor cells attract and dynamically interact with monocytes/macrophages. To find biomarkers of disease progression in human melanoma, we used unbiased RNA sequencing and secretome analyses of tumor-macrophage co-cultures. Pathway analysis of genes differentially modulated in human macrophages exposed to melanoma cells revealed a general upregulation of inflammatory hallmark gene sets, particularly chemokines. A selective group of chemokines, including CCL20, CCL15 and CCL8, was actively secreted upon melanoma-macrophage co-culture. Because we previously described the role of CCL20 in melanoma, we focused our study in CCL8 and CCL15, and confirmed that in vitro both chemokines contributed to melanoma survival, proliferation and 3D invasion, through CCR1 signaling. In vivo, both chemokines enhanced primary tumor growth, spontaneous lung metastasis and circulating tumor cell (CTC) survival and lung colonization in mouse xenograft models. Finally, we explored the clinical significance of CCL8 and CCL15 expression in human skin melanoma, screening a collection of 67 primary melanoma samples, by multicolor staining and quantitative image analysis of chemokine-chemokine receptor content at the single cell level. Primary skin melanomas displayed high CCR1 expression, but there was no difference in its level of expression between metastatic and non-metastatic cases. By contrast, the comparative analysis between these two clinically divergent groups showed a highly significant difference in the cancer cell content of CCL8 (P= 0.025) and CCL15 (P< 0.0001). Kaplan–Meier curves showed that high content of CCL8 or CCL15 in cancer cells correlated with shorter disease-free and overall survival (log-rank test, p< 0.001). Our results highlight the role of CCL8 and CCL15, which are highly induced by melanoma-macrophage interactions in biologically aggressive primary melanomas, and could be clinically applicable biomarkers for patient profiling.

## Introduction

Melanoma is a form of skin cancer characterized by its ability to metastasize to distant organs [1]. Identifying early-stage cutaneous melanoma patients that are likely to develop distant metastasis is clinically relevant because those patients may benefit from immunotherapy [2]. There is a need to develop prognostic markers to identify patients at higher risk of metastatic disease as well as predictive markers of response to immunotherapy, to guide therapeutic decisions and clinical trials with precision. Current models of metastatic progression propose that cancer cell invasion and metastasis are strongly influenced by contextual signals emanating from the stroma of the primary tumor; therefore, analysis of tumor-stroma interactions may be a source of new prognostic markers [3].

Important molecular partners underlying tumor-stroma interactions are secreted proteins of the chemokine family, acting through their cell receptors to guide migration and positioning of infiltrating cells in the tumor microenvironment [4]. Chemokines may act on nearby cells (paracrine action) as well as on the cells that secrete them (autocrine action) and may have both anti-tumoral and pro-tumoral functions. The chemokine milieu and the particular chemokine receptors expressed by tumor and immune cells are tumor type specific and strongly heterogeneous. Melanoma, in particular, is an inflammatory cancer type very influenced by the chemokine network [5, 6]. Some chemokines were originally identified in melanoma for their autocrine growth promoting function, such as *melanoma growth stimulating activity* (MGSA, currently known as CXCL1) [7], or IL-8 (known as CXCL8) which acts as an autocrine/paracrine growth factor activating CXCR1 and CXCR2 [8]. Several chemokine paracrine loops are involved in the response to ultraviolet radiation during melanomagenesis, where CCL2/CCL7 secreted by melanocytes recruit CCR2+ immune cells to initiate the process [9].We described the CCL20/CCR6 axis, in which CCL20-secreting macrophages promoted tumor survival, growth and invasion through CCR6+ melanoma cells [10]. Importantly, the stromal content of CCL20 was associated with patient survival in primary melanoma [11].

Among the most abundant cells recruited and modified by tumors are tumor associated macrophages (TAMs), which display a spectrum of states associated with diverse functions such as angiogenesis, immunosuppression, inflammation or extracellular matrix remodeling [12]. Diversity of TAMs is further supported by their origin from either blood monocytes or tissue resident macrophages or by their response to stimuli provided by different types of tumor cells or other infiltrating immune cells [13]. Macrophage phenotype and function were simplified by the M1/M2 in vitro classification, which assumed that TAMs were preferentially M2 biased in advanced solid tumors [14]. More recently, in vivo single-cell analysis has revealed an unsuspected heterogeneity of TAM phenotypes in various types of human cancer including melanoma [15]. Our hypothesis is that understanding the complexity of TAM-tumor cell crosstalk, may be a source of prognostic markers and immunotherapy targets in human melanoma. Our previous work showed that primary tumors from patients that developed metastasis during follow-up are characterized by TAMs with a cytokine secretory phenotype, including CCL20, TNF and VEGFA [11]. We also showed that a particular TAM oncogenic activation is mediated by Activin A and regulated by p53 and NFκB co-activation [16]. To continue exploring TAM-tumor cell crosstalk in human melanoma, we now used RNA sequencing and secretome analyses to identify the cytokine and chemokine milieu during TAM programing by melanoma cells.

## Materials and Methods

### Monocyte Isolation and Cell Culture

Peripheral blood mononuclear cells (PBMCs) were isolated from buffy-coats from healthy donors over a ficoll gradient. Monocytes were purified by magnetic cell sorting using anti-CD14 tagged microbeads (Miltenyi Biotech, Bergisch Gladbach, Germany) and cultured at 0.5 ×106/ml for 7 days containing GM-CSF (10 ng/ml, Immunotools, Friesoythe, Germany) or M-CSF (10 ng/ml, Immunotools) to generate M1 or M2 macrophages, respectively. All cells, including melanoma cell lines BLM, A375 and Skmel-103 were cultured in RMPI-1640 medium (Gibco, Waltham, MA, US) supplemented with 10% fetal calf serum (FCS, Sigma-Aldrich, St. Louis, MO).

### In vitro measurements

Melanoma cells were cocultured with M1 or M2 macrophages at 1:2 ratio (melanoma:macrophage) for 24 or 72 hours in order to perform transcriptomic or secretomic analyses, respectively. Conditioned media were collected for multiplex ELISA arrays (Quantibody, RayBiotech, GA) and for CCL8 (PeproTech Inc, Waltham, MA, US) and CCL15 (Abcam, Cambridge, UK) ELISA. For transcriptomic analyses, conditioned macrophages were separated from melanoma cells with CD14-microbeads. For RNAseq, total RNA was isolated from three independent preparations and processed at BGI (https://www.bgitechsolutions.com), using the DNBseq-G400 platform. Differential gene expression was assessed by using DEseq2 algorithms using the parameters Fold change >2 and adjusted p value <0.05. For GSEA, different hallmark gene sets available at the website (http://www.broad.mit.edu/gsea/) [17, 18] were used, as well as the gene sets “GOMF_CHEMOKINE_ACTIVITY” and “KEGG_CYTOKINE_CYTOKINE RECEPTOR_INTERACTION”. Data were deposited in NCBI’s Gene Expression Omnibus and are accessible through GEO Series accession numbers GSE171277 and GSE242674. Real-time PCR (qPCR), oligonucleotides (supplementary material, Table S1) were designed with Roche software. RNA was extracted (NucleoSpin RNA-purification kit, Macherey-Nagel Dueren, Germany) and retrotranscribed cDNA quantified using the Universal Human Probe Roche library (Roche-Diagnostics, Barcelona, Spain). Assays were made in triplicate and normalized to *TBP* expression (ΔΔCT method). For expression analysis of chemokine receptors, whole-cell lysates were subjected to western blotting with the indicated antibodies (supplementary material, Table S1).

### 3D-invasion assay

BLM spheroids were produced by seeding 104 cells per well in a 96-well U-bottom plate (Thermofisher Scientific, Waltham, MA, US) for 5-7 days and embedded into 2 mg/ml collagen type-I (StemCell Technologies, Vancouver, Canada) gels. Serum-deprived media ±recombinant human CCL8 or CCL15 (R&D Systems, Minnneapolis, MN) and ±5 μg/ml neutralizing antibodies for CCR1, CCR3 and CCR5 (supplementary material, Table S1) were added on top of the gels, and spheroids were allowed to invade for 72 hours. For monocyte invasion, 5x104 cells were cultured in 0.5% FCS medium and added on top of collagen gels containing chemokines and allowed to invade for 48 hours. For quantification, cells were stained with propidium iodide (Sigma-Aldrich) and imaged with the ACS_APO 10x/NA 0.30 objective of an inverted confocal microscope (Leica, SPE, Wetzlar, Germany). Spheroids were imaged at central planes and maximum invading distances from the spheroid edge were assessed at multiple random directions. Percentage of invading monocytes (>30 μm) were assessed through 200 μm depth *z*-stacks. Invasion was quantified using ImageJ2 software [19].

### Survival and proliferation assays

For cell viability assays, melanoma cells or monocytes were cultured in serum-deprived media ±chemokines for 5 and 2 days. Media were then removed and cells fixed with 4% formaldehyde for colorimetric crystal violet assay (Thermofisher Scientific). For proliferation assays, melanoma cells were cultured in serum-deprived medium for 24 hours ±chemokines, adding 10 μg/ml 5-bromo-2′-deoxyuridine (BrdU, Sigma-Aldrich) 1 hour before fixation. Cells were permeabilized and stained with 5 μg/ml anti-BrdU (R&D Systems). Samples were imaged with a glycerol-immersion ACS_APO 20x/NA 0.60 objective (Leica, SPE), DAPI-stained nuclei were segmented, and BrdU Mean Fluorescence Intensity (MFI) assessed with the ImageJ2 [10, 16].

### In vivo human melanoma models

NOD/Scid/Il2rg-/- (NSG) mice (The Jackson Laboratory, Bar Harbor, ME) were maintained under specific pathogen-free conditions. At age of 6 weeks (∼23 grams), male or female mice were subcutaneously inoculated with 106 melanoma cells suspended in 100 μl PBS ±200 ng/ml chemokines. Mice were then subcutaneously injected with PBS ±1 μg human chemokines every 3 days, and primary tumor volumes (width^2 x length, cm3) measured 17-19 or 21-23 days post-injection for BLM or A375 models, respectively. When xenografts were 1 cm width, BLM-bearing mice were sacrificed and lungs resected for histological quantification of micrometastases. Some mice were injected intraperitoneally 1 mg BrdU 4h before sacrifice and 20 hours after the last chemokine doses. For lung colonization experiments, male mice were subcutaneously injected with PBS ±1 μg chemokines 15 minutes before tail intravenous injection of 0.5×106 melanoma cells suspended in 100 μl PBS ±200 ng/ml chemokines. Two additional doses of chemokines were subcutaneously injected every 2 days. At day 7th, lungs were extracted for histologic analyses. Procedures were approved by the IiSGM animal care/use and Comunidad de Madrid committees (Proex: 084/18 and 296.4/22).

### Study Cohort and Selection Criteria

Patient samples were collected following the approval of Gregorio Marañón Hospital ethics committee and informant consent was obtained for each patient. A formalin-fixed and paraffin-embedded (FFPE) primary cutaneous melanoma cohort of 67 patients was used, with a >2 mm Breslow thickness and a median follow-up of 81 months, excised between 1998 and 2015 in our institution. This cohort includes 31 samples from patients who were disease-free for at least 10 years of follow-up (non-metastasizing primary melanomas) and 36 clinically aggressive samples developing distant metastasis (metastasizing primary melanomas, with 23/36 melanoma-related deaths). Pathological AJCC staging II-IV assessment was obtained by sentinel lymph node biopsy and distant metastasis evaluation by computed tomography at the time of diagnosis. Metastasizing and non-metastasizing primary tumors had comparable Breslow thickness (Mann– Whitney, p= 0.37). Five patients at stage IV were excluded from disease-free survival (DFS) analysis, but not from overall survival (OS).

### Multicolor Fluorescence Confocal Microscopy

FFPE sections were deparaffinized, rehydrated, and unmasked by steaming in 10 mM sodium citrate buffer pH 9.0 (Dako Glostrup, Denmark) for 7 minutes. Slides were blocked with 5 μg/ml human immunoglobulins solved in blocking serum-free medium (Dako) for 30 minutes, and then sequentially incubated with 5–10 μg/ml primary antibodies, overnight, at 4°C, and proper fluorescent secondary antibodies (Jackson Immunoresearch, West Grove, PA, US) for 1h. Washes were performed in PBS containing 0.05% Tween-20. Single-cell quantification was performed in 3–5 20× fields. MFI of proteins of interest was obtained at manually depicted tumor cell nests or at segmented CD68+ TAM using the ‘analyze particle’ plugging of ImageJ2 software (11) (12) (17). For mouse tumors, BrdU MFI of DAPI-stained segmented nuclei for proliferation assays; or quantification of NG2+ lung micrometastases/colonies. Samples were imaged with ACS_APO 20x/NA 0.60 objective (Leica, SPE).

### Statistical Analyses

Kaplan-Meier curves were used to analyze the correlation with patient disease-free and overall survival, using Youden’s index to determine the cutoff point equally specific and sensitive. Cox-regression method (univariate and multivariate) was used to identify independent prognostic variables and Mann-Whitney test to evaluate the association with clinicopathological features. Paired and unpaired t-test and Log-rank analyses have also been used in this study (Graph-Pad software, San Diego, CA, United States), as indicated. p <0.05 was considered statistically significant.

## Results

### Chemokine upregulation by melanoma-macrophage interactions

As a source of biomarkers for human melanoma progression, we used RNA-sequencing data and secretome panel validation of melanoma-macrophage co-cultures. To consider the wide spectrum of macrophage polarization, we differentiated human macrophages from blood monocytes in the presence of either GM-CSF or M-CSF, which are representative of the M1 and M2 states, respectively (GM-CSF for M1 pro-inflammatory and M-CSF for M2 anti-inflammatory macrophages) [20]. The melanoma cell line BLM was co-cultured with either M1 or M2 macrophages and then separated for RNAseq; whereas conditioned media were collected for secretome panel examination. Thus, tumor–exposed macrophages (TEM), either M1 or M2, and macrophage-exposed melanoma cells, BLM(M1) or BLM(M2), were compared to unexposed ones for Gene Set Enrichment Analysis (GSEA) (Figure 1A). The analysis showed a general upregulation of inflammatory hallmark gene sets, being the highest upregulated pathway ‘TNF_signaling_via_NFκB’ in all the co-cultured cells, including TEM1 and TEM2, irrespective of their initial polarization state, as well as BLM tumor cells exposed to either M1 or M2 macrophages. Specific cytokine and chemokine inflammatory gene sets were additionally interrogated to deepen on the inflammatory response upregulation. Consistently, both chemokine activity and cytokine-cytokine receptor interaction were highly upregulated in all the co-cultured cells, compared to unexposed ones. Conditioned media from the M1/M2 macrophage-melanoma co-cultures were subjected to secretome analysis and compared to unexposed cell supernatants (Figure 1B, C; supplementary material, Table S2). A larger set of chemokines was induced in the M2-melanoma co-cultures, compared to the M1-melanoma ones, which up-regulated only a selective group of chemokines. Interestingly, only three chemokines, CCL8, CCL15 and CCL20, were highly induced in both M1 and M2 macrophage-melanoma co-cultures. These data indicated that melanoma-macrophage interactions differentially induced a particular group of chemokines, regardless of the initial M1/M2 polarization state of the macrophages. Because we already described the oncogenic potential of CCL20 in melanoma [10, 11], we focused our study in CCL8 and CCL15, and confirmed by ELISA the induction of both chemokines in melanoma-macrophage co-cultures (Figure 1D). To search for the cell source of these chemokines, we performed qPCR analysis of separated macrophages and tumor cells after co-culture, showing high induction of *CCL8* and *CCL15* mRNA in both TEM1 and TEM2 (Figure 1E). Besides, BLM cells expressed more *CCL15* mRNA in response to M1 than to M2, whereas BLM expressed more *CCL8* mRNA in response to M2 than to M1.

**Figure 1.**
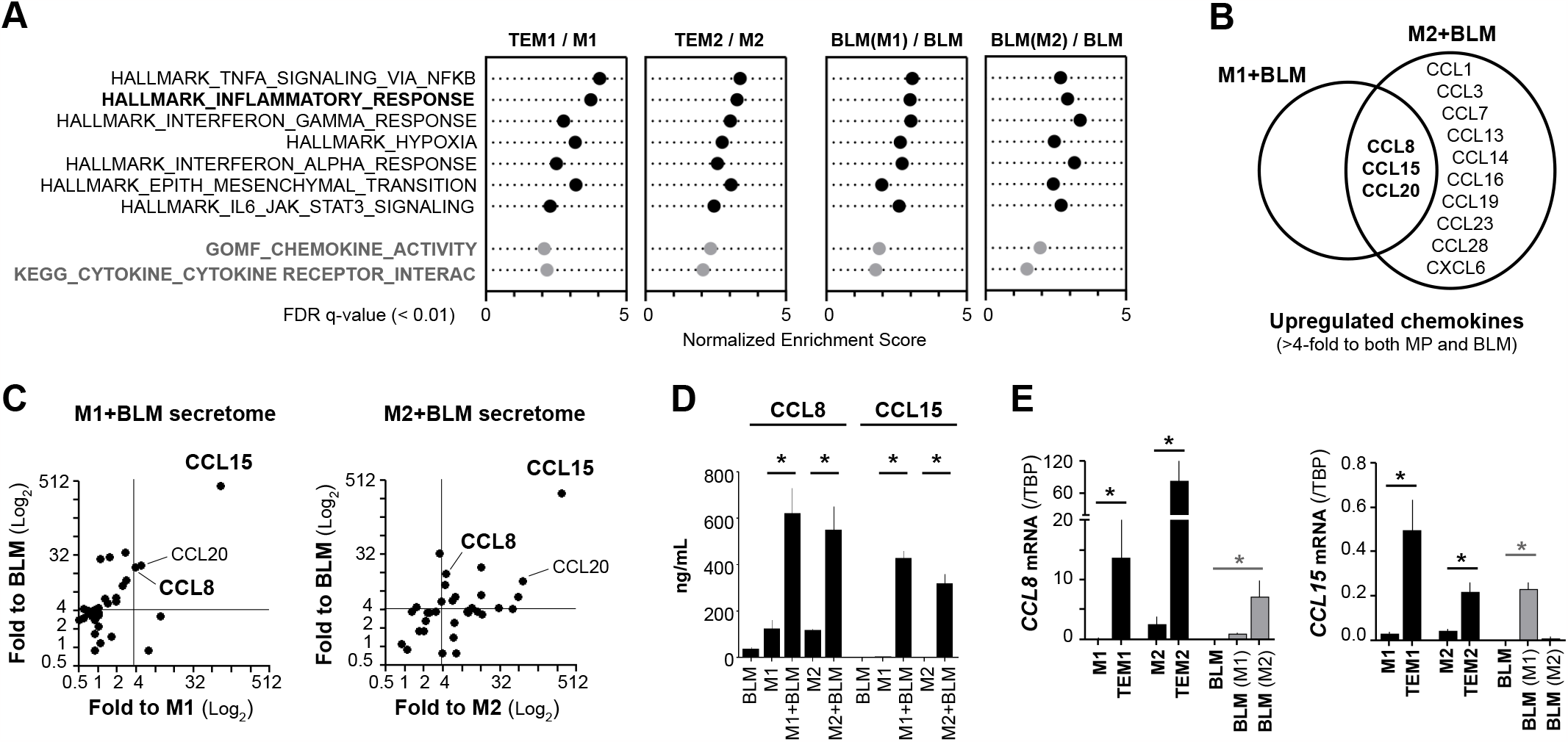
Co-culture of macrophages and melanoma cells induces the expression of CCL15 and CCL8 chemokines. M1 and M2 macrophages were co-cultured with BLM melanoma cells for transcriptomic (24 h) and secretomic (72 h) analyses. (A) Gene set enrichment analyses (GSEA) comparing 24 hours exposed vs un-exposed isolated cells, as indicated (n=3 donors). Normalized Enrichment Score (NES) values from the main upregulated hallmark_pathwaysfor all groups,are shown. GOMF and KEGG gene sets were additionally studied for cytokine/ chemokine inflammatory pathways. False Discovery Rates are represented (FDR q value <0.01). (B) Representative scheme highlighting the coincident most upregulated secreted chemokines (fold >4 to both unex-posed macrophages and melanoma cells). (C) Secretomic analysis (38 chemokines) of 72 hours co-cultured supernatants comparing M1+ BLM or M2+BLM with unexposed macrophages (x-axis) or BLM (y-axis) cells. Fold logarithmic average changes are shown (n=4 donors). (D) CCL8 and CCL15 secretion assessed at 72 h macrophages ±BLM conditioned media. Mean ±standard deviation (SD) values are shown (n= 5 do-nors). (E) CCL8 and CCL15 mRNA expression levels in mutually conditioned isolated cells. Mean ±SD values relative to TBP mRNA are shown (n= 4). Significant differences to the respective untreated controls, are shown (*p< 0.05, unpaired t-test).

### CCL8 and CCL15 induce melanoma growth and invasion through CCR1 signaling

Chemokines can regulate tumor progression by acting directly on the tumor cells, supporting tumor cell survival, proliferation and invasion. We used serum deprivation to test cell viability and proliferation of three different melanoma cell lines, in the absence or in the presence of increasing doses of CCL8 or CCL15, showing that both chemokines induced cell survival and proliferation (Figure 2A, B). For 3D invasion assays, BLM melanoma spheroids were embedded in collagen matrices in the absence or in the presence of CCL8 or CCL15, showing dose–response 3D invasion in response to both chemokines (Figure 2C). Because human CCL8 interacts with multiple chemokine receptors, including CCR1 and CCR5, and CCL15 binds to CCR1 and CCR3, we used blocking antibodies to these receptors to show that CCR1 was the primary signaling receptor involved in melanoma 3D invasion. Furthermore, CCR1 was highly expressed by western blotting in cell extracts from the three-melanoma cell lines included in the functional assays (Figure 2D). These results indicated that CCR1 is the primary signaling receptor involved in the direct functional response of melanoma cells to both CCL8 and CCL15 chemokines. An equivalent molecular weight band corresponding to CCR1 was detected on peripheral blood monocytes, included as controls, which also responded to both chemokines in functional assays (Figure 2D, E). These results suggest that CCL8 and CCL15 chemokines may contribute to melanoma progression by direct effects on tumor cells expressing CCR1, as well as, indirectly by recruiting monocytes/macrophages towards the tumor microenvironment.

**Figure 2.**
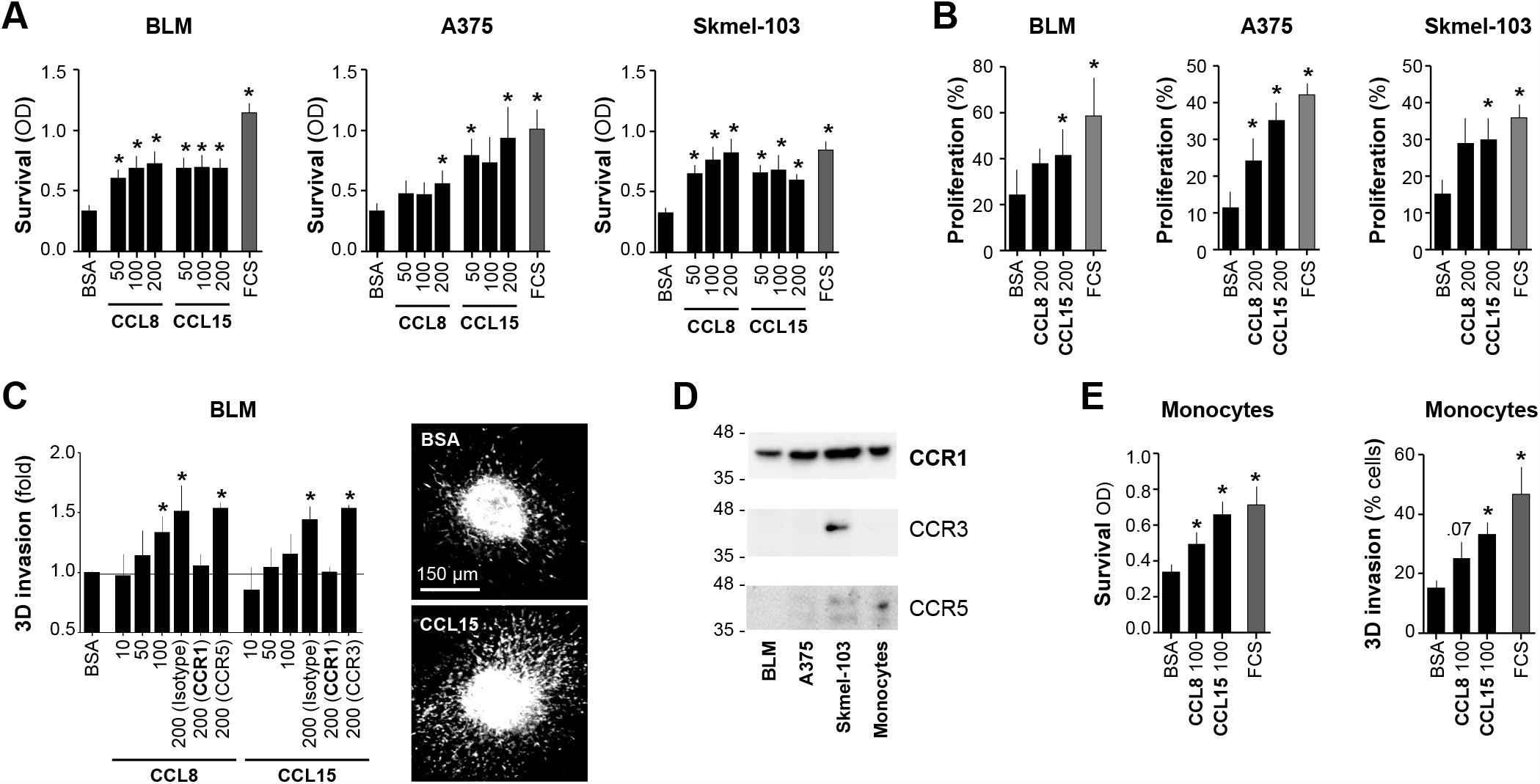
Exogenous CCL8 and CCL15 chemokines induce survival, invasion and proliferation of melanoma cells in vitro. (A) Serum deprivation survival assays of BLM, A375 and Skmel-103 melanoma cell lines in response to increasing concentra-tions of CCL8 and CCL15 chemokines (ng/ml), and to FCS 1%, as positive control. Mean ±SD optical density (OD) values, are shown (n= 4). (B) Proliferation response of melanoma cell lines to 200 ng/ml exogenous CCL8 and CCL15, and to FCS 1% as positive control. Mean ±SD percentages of BrdU+ nuclei, are shown (n= 3). (C) BLM spheroids embedded in 3D-collagen and allowed to invade for 72 h in the presence of increasing concentrations of exogenous CCL8 and CCL15, as indicated (ng/ml). Functional chemotactic axes were identified by using neutralizing antibodies against CCR1, CCR3, CCR5 and their corresponding control isotypes (5 μg/ml). Mean ±SD fold-invasion values, are shown (n= 3-6). Images show representative collagen-embedded spheroids showing basal and CCL15 stimulated invasion. Scale bar, 150 μm. (D) Immunoblots showing CCR1, CCR3 and CCR5 expression in whole-lysates of three melanoma cell lines and isolated CD14+ monocytes. (E) Serum deprivation survival and 3D-collagen invasion assays of monocytes in response to 100 ng/ml exogenous chemokines, including FCS 1%, as positive control (n= 5 donors). Significant differences to the respective untreated controls, are shown (*p< 0.05, paired t-test).

### CCL8 and CCL15 support melanoma progression in vivo

To explore the role of CCL8 and CCL15 in tumor progression in vivo, we injected subcutaneously two melanoma cell lines in NSG mice and measured primary tumor growth and spontaneous lung colonization, in the absence or presence of each chemokine (Figure 3A). Both chemokines enhanced primary tumor size and proliferating cells, stained by BrdU incorporation into BLM tumors in vivo (Figure 3B; supplementary material, Figure S1A). Because primary tumor growth is uncoupled to metastatic dissemination, we also quantified spontaneous lung metastasis by the highly invasive BLM cells and found that both chemokines enhanced the number and size of metastatic foci (Figure 3C; supplementary material, Figure S1B). A critical step in the metastatic process is survival of circulating tumor cells (CTC) and colonization of hostile distant organs, therefore, we also tested the effect of CCL8 and CCL15 in lung colonization assays, in which BLM melanoma cell line was injected intravenously (i.v.) to mimic CTC. Direct i.v. injection of BLM cells in the presence of both chemokines induced more lung tumor colonies, suggesting a role for CCL8 and CCL15 in facilitating survival and colonization of CTC (Figure 3D). Therefore, both chemokines enhanced primary tumor growth, spontaneous lung metastasis and CTC survival and lung colonization.

**Figure 3.**
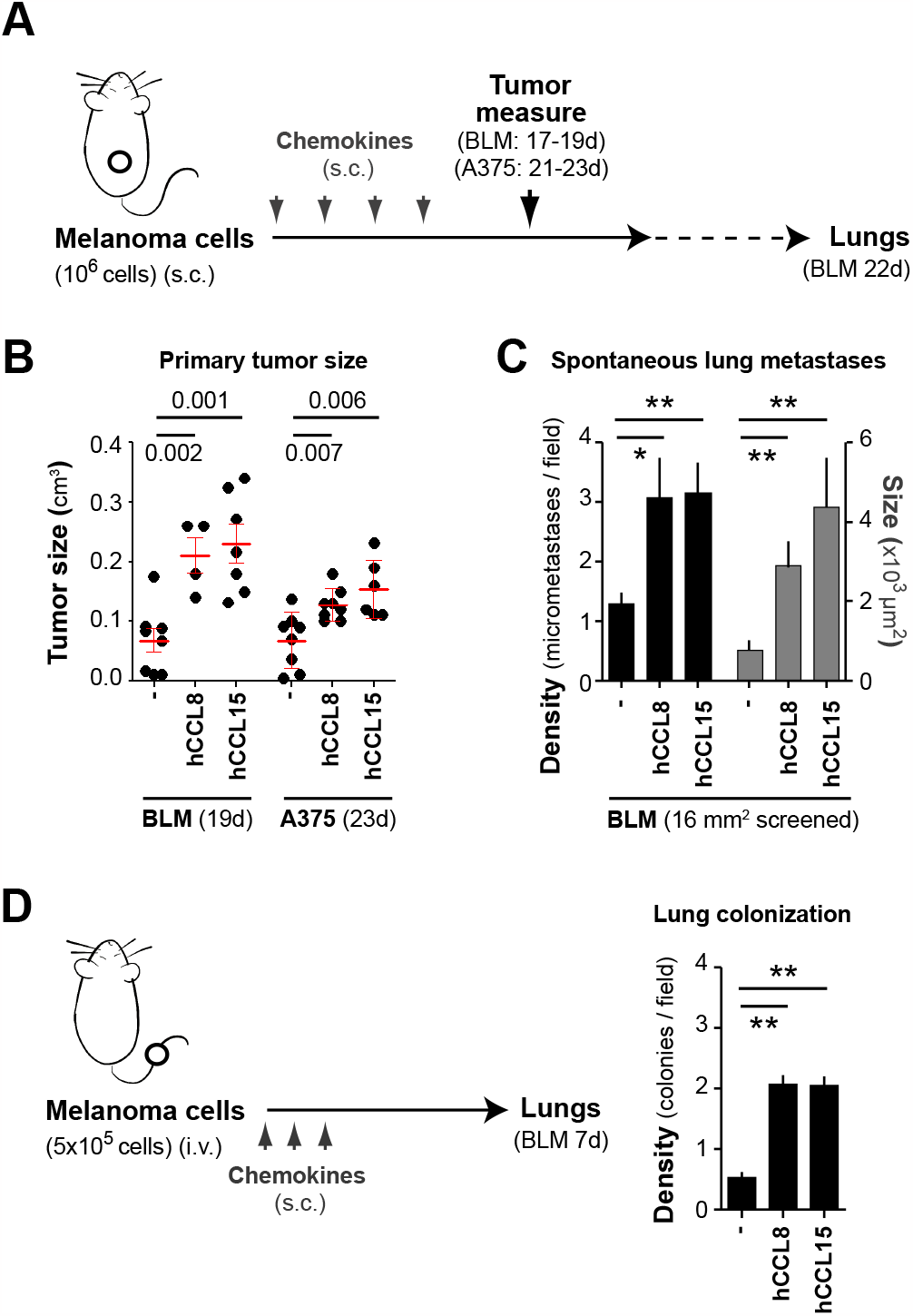
Exogenous CCL8 and CCL15 induce human melanoma tumor progression and lung colonization in vivo. (A) Schematic representation of our model of human melanoma xenografts development in immune-compromised NSG mice. Plot showing A375 and BLM primary tumor volumes at indicated times (cm3), including the mean ± standard error of the mean (SEM) (n= 4-8 mice). (C) Mean ±SD density (nº/field) and size (μm2) values of BLM spontaneous micrometastases, are shown (n= 3-7 mice). (D) Schematic representation of our model of lung colonization assays and quantification of BLM lung colonies at 7th day post-tail injection, quantified as in C (n= 3). Significant differences to the respective untreated controls are shown at C and D panels (*p< 0.05, unpaired t-test), or indicated at B.

### CCL8 and CCL15 expression by primary human melanoma tissues

Our RNAseq and secretome screenings identified CCL8 and CCL15 as key chemokines induced during melanoma‐myeloid crosstalk; furthermore, both chemokines enhanced primary tumor growth and spontaneous lung metastasis in mouse models. Next, we explored the clinical significance of their expression in the microenvironment of human skin melanoma in a collection of 67 primary melanoma samples with clinical information and survival status of the patients (supplementary material, Table S3). Primary melanomas were classified as non-metastasizing or metastasizing, regarding subsequent development of metastasis during patient follow-up for 10 years. To allow quantification of the chemokine-chemokine receptor content within CD68+ macrophages or melanoma cells, we set-up conditions for multicolor staining of CCL8, CCL15, CCR1, CCR3 and CCR5, and compared the relative expression data in segmented cells with clinicopathological factors (supplementary material, Table S3). Regarding quantification within CD68+ TAMs, the comparative analysis of non-metastasizing versus metastasizing primary tumors showed low expression of CCR5 and no mayor differences in CCL8, CCL15 or CCR1 content, whereas CCR3 MFI was higher in TAMs from clinically aggressive melanomas (Figure 4A, C; supplementary material, Table S3). Single-cell measurements for individual macrophages in non-metastasizing and metastasizing cases of CCR1, CCR3 and CCR5 showed a subpopulation of TAMs highly expressing CCR3 in metastasizing samples (supplementary material, Figure S1C). Thus, the association of high CCR3 expression by TAMs with aggressive melanomas suggests that this receptor may promote their tumor supportive functions. Regarding quantification within tumor cells (TCs), the comparative analysis between non-metastasizing and metastasizing primary tumors showed a highly significant difference in the cancer cell content of CCL15 (P< 0.0001), with lower significance in the case of CCL8 expression (P= 0.025) (Figure 4B, C; supplementary material, Table S3). Concerning chemokine receptor expression, cancer cells barely expressed CCR3 or CCR5 and highly expressed CCR1, but with no differences between non-metastasizing and metastasizing primary tumors. Next, we used the Kaplan–Meier method to assess the clinical relevance of quantifying chemokine-chemokine receptor content, by TCs or TAMs, in melanoma tissues from primary melanoma patients (Figure 5A). Patients were stratified as ‘high’ or ‘low’, using the best cutoff value extrapolated from ROC (receiver operating characteristic) curve to calculate 10-years disease-free survival and overall survival curves. Compared to parameters measured in TAMs, that were borderline or less significant in Kaplan–Meier curves, differences in the cancer cell content of CCL8 or CCL15 were highly significant, showing that high CCL8 or CCL15 content in TCs correlated with shorter DFS and OS (log-rank test, p< 0.001). Because CCL15 expression by melanoma cells was higher than CCL8 (Figure 4B), we analyzed whether CCL15 quantification was an independent prognostic factor, in a multivariate regression analysis including gender, age, location, histologic type, ulceration, Breslow and stage parameters (Table 1). The regression analysis showed that CCL15 expression at cancer cells was an independent prognostic factor for DFS (p< 0.001) and OS (p= 0.015) in this cohort. Our results with patient samples highlight the role of CCL15 expression by cancer cells in biologically aggressive primary melanomas, suggesting that it could be a clinically applicable biomarker for patient profiling. Altogether, our data indicate that CCL8 and CCL15, highly induced by melanoma-macrophage interactions under certain conditions, may be signaling through CCR1 in melanoma cancer cells to drive melanoma progression (Figure 5B).

**Table 1.** Univariate and multivariate Cox regression analyses for 10-year disease-free and overall survival. Abbreviations: a.u., arbitrary units; CI, confidence interval; F, female; H/L, head/limb; HR, hazard ratio; M, male; MFI, mean fluorescence intensity; nod, nodular; T, trunk; TAM, tumor-associated macrophage; TC, tumor cell. MFI cutoff points are indicated.

| Univariate | Disease-free survival |  |  | Overall survival |  |  |
| --- | --- | --- | --- | --- | --- | --- |
|  | HR | 95% CI | P | HR | 95% CI | P |
| Gender (F vs M) | 1.687 | 0.93-3.03 | 0.081 | 0.966 | 0.48-1.92 | 0.913 |
| Age (years) | 1.003 | 0.98-1.02 | 0.774 | 1.001 | 0.52-2.14 | 0.874 |
| Location (H/L vs T) | 1.564 | 0.79-3.08 | 0.196 | 1.059 | 0.43-1.92 | 0.812 |
| Subtype (nod vs others) | 0.935 | 0.51-1.71 | 0.827 | 1.165 | 0.58-2.33 | 0.666 |
| Ulceration (yes vs no) | 1.298 | 0.71-2.36 | 0.395 | 1.937 | 0.97-3.82 | 0.057 |
| Breslow (mm) | 1.108 | 1.04-1.17 | 0.001 | 1.09 | 1.01-1.17 | 0.023 |
| Stage (II vs III-IV) | 1.421 | 0.73-2.74 | 0.296 | 2.184 | 1.04-4.56 | 0.038 |
| TAM CCR3 (MFI, >78 a.u.) | 2.793 | 1.23-6.31 | 0.014 | 2.285 | 0.90-5.77 | 0.081 |
| TC CCL8 (MFI, >80 a.u.) | 2.359 | 1.12-4.95 | 0.024 | 7.893 | 2.25-27.6 | 0.001 |
| TAM CCL15 (MFI, >69 a.u.) | 2.283 | 1.06-4.88 | 0.033 | 2.838 | 1.13-7.08 | 0.025 |
| TC CCL15 (MFI, >98 a.u.) | 7.923 | 3.37-18.6 | <0.001 | 4.629 | 1.83-11.7 | 0.001 |
| Multivariate | HR | 95% CI | P | HR | 95% CI | P |
| Gender (F vs M) | 1.51 | 0.62-3.39 | 0.337 | 0.685 | 0.24-1.92 | 0.474 |
| Age (years) | 1.015 | 0.98-1.04 | 0.331 | 1.005 | 0.96-1.04 | 0.803 |
| Breslow (mm) | 1.092 | 1.00-1.19 | 0.048 | 1.057 | 0.96-1.16 | 0.244 |
| Stage (II vs III-IV) | 1.038 | 0.45-2.35 | 0.929 | 2.291 | 0.96-5.46 | 0.061 |
| TC CCL15 (MFI, >98 a.u.) | 6.449 | 2.55-16.2 | <0.001 | 3.303 | 1.25-8.66 | 0.015 |

**Figure 4.**
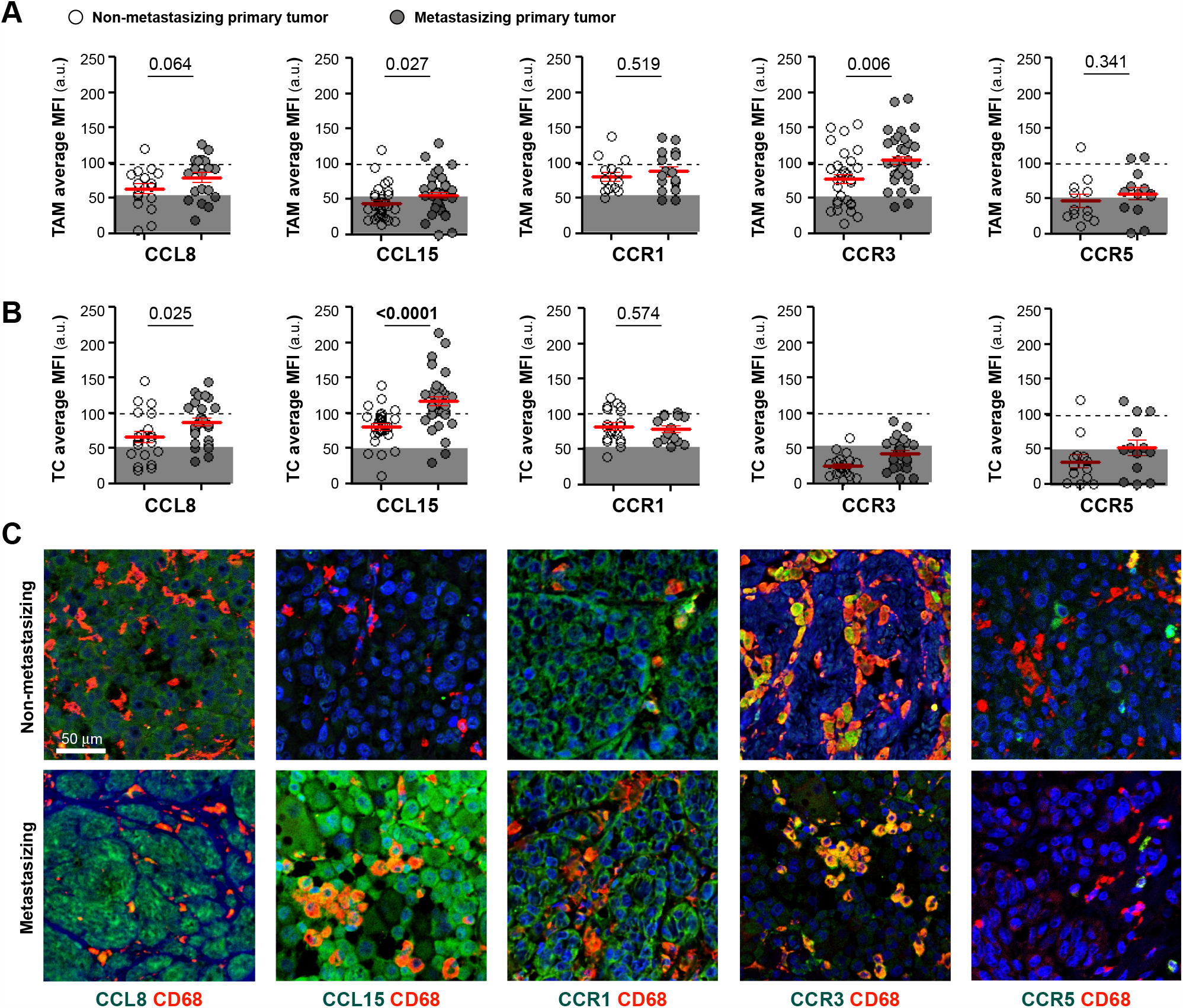
Chemokines expression in human primary melanoma and its correlation with patient survival. TAM (A) and tumor cell (B) average MFI of CCL15, CCL8, CCR1, CCR3 and CCR5 in non-metastasizing and metastasizing primary tumors.Mann-Whitney statistical analysis was used to compare non-metastasizing vs metastasizing melanomas; p-values are shown.(C) FFPE human melanoma samples stained for CD68 (TAM marker, red) and CCL15, CCL8, CCR1, CCR3 or CCR5 (green). Scale bar, 50 μm.

**Figure 5.**
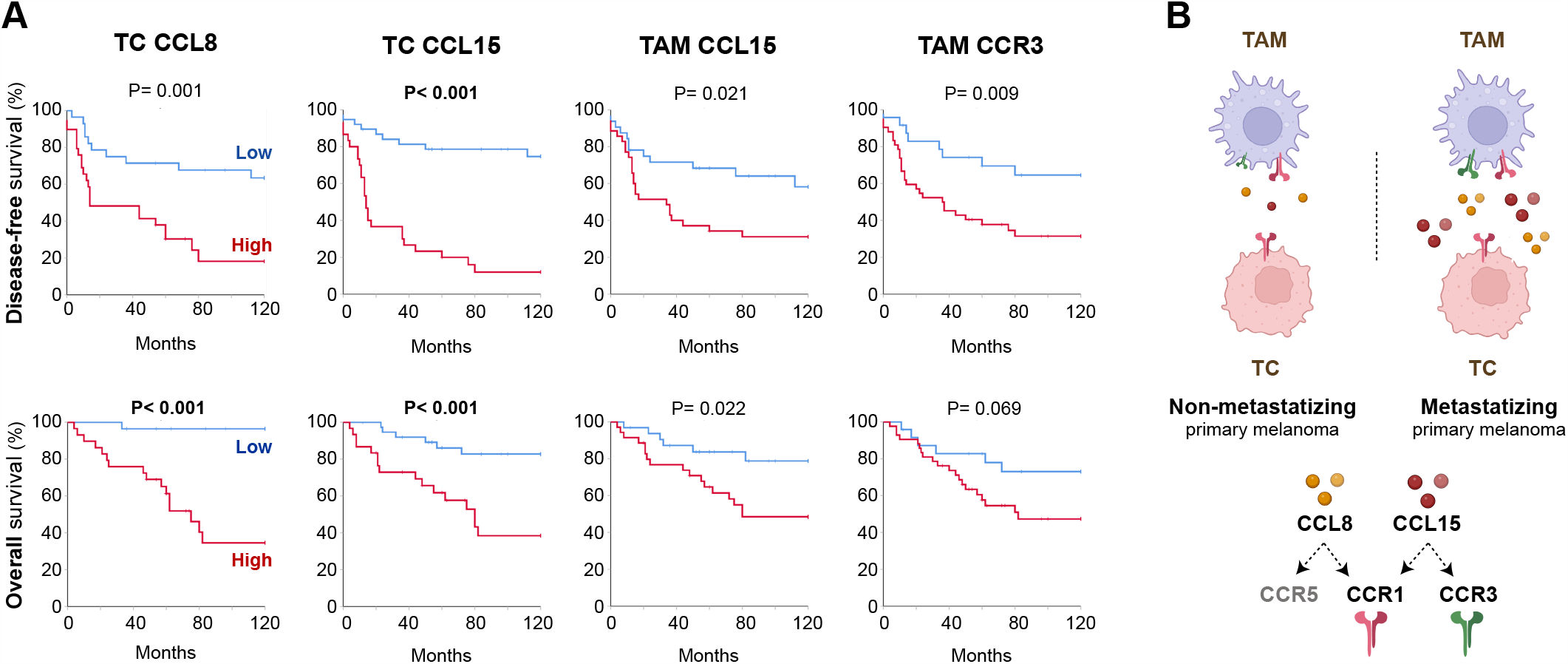
Kaplan–Meier curves and summarizing schematic model. (A)Disease-free and overall survival 10-year Kaplan–Meier curves. Youden’s index was used to choose a cutoff point to classify the 67 primary melanomas as ‘low’ or ‘high’: TC CCL8 (MFI, 80 a.u.), TC CCL15 (98a.u.),TAM CCL15 (69 a.u.) and TAM CCR3 (78 a.u.). Log-rank p values, are shown. (B) Summarizing model, created with BioRender.com
.

## Discussion

At present, we have limited understanding of the complexity of TAM-tumor cell crosstalk in human cancer. To find biomarkers of disease progression and prognosis in melanoma patients, we explored up-regulated secreted factors induced by melanoma-macrophage interactions. We previously used this approach to find that Activin A, a cytokine of the TGF-family secreted by both melanoma cells and macrophages, was involved in the protumoral deviation of TAM [16]. Indeed, INHBA which is the gen coding for Activin A, is nowadays recognized as a marker of pro-angiogenic INHBA+ TAMs in single-cell transcriptional atlas across cancer types [15, 21]. Now, we focused our attention on up-regulated chemokines, finding CCL8, CCL15 and CCL20 as key chemokines mediating TAM-tumor cell crosstalk in human melanoma. We previously described the paracrine CCL20/CCR6 axis in human melanoma [10], and we now describe the autocrine/paracrine role of CCL8 and CCL15 through CCR1. We show that high CCL8 or CCL15 protein expression by cancer cells was associated with bad prognosis and metastasis development in a cohort of 67 skin melanoma patients. CCL15 protein expression was an independent prognostic factor in regression analysis suggesting that it may be a useful biomarker for the prognosis of patients with primary cutaneous melanoma.

A complex network of intercellular communication in the TME is mediated by chemokines, which are chemotactic cytokines actively secreted by both cancer and non-cancer cells [22]. Chemokines attract and position non-cancer cells expressing the corresponding chemokine receptors; these recruited cells also secrete more chemokines into cancerous tissues. Thus, chemokines are key molecular players governing dynamic changes in the TME, but they are cancer-type specific, context dependent and a systematic understanding of the rules that govern chemokine mediated cell-to-cell communication at the global tissue-level is still lacking. Although melanoma is one of the tumors where the role of chemokines and their receptors has been extensively investigated, the role of CCL15 was not known and contradictory data were reported for CCL8 chemokine in human melanoma. Transcriptome data from the Cancer Genome Atlas, analyzed by two groups, showed that skin melanoma tissues expressing high levels of CCL8 had better patient prognosis [23, 24]. By contrast, increased *Ccl12* mRNA expression, the mouse orthologous of human CCL8, was identified in a metastasis forming primary melanoma mouse model, although no statistical difference between metastatic and non-metastatic groups from a cohort of primary melanoma patients was found for *CCL8* RNA expression [25]. Our data show an association of high CCL8 protein expression by melanoma cells with worse patient prognosis, which is in line with data from other cancer types, as breast cancer, in which CCL8 expression is associated with worse patient prognosis [26]. Indeed, CCL8, also known as monocyte chemoattractant protein 2 (MCP-2), is mainly involved in monocyte recruitment, which is generally associated with progressing tumors [27].

Cancer cells express multiple chemokine receptors responding directly to chemokines in the TME. In agreement with previous results [28], we detected high CCR1 expression by melanoma cell lines, which directly responded to CCL8 and CCL15 chemokines in survival, proliferation and 3D invasion assays. Primary skin melanomas also displayed high CCR1 expression, but there was no difference in the level of protein expression between metastatic and non-metastatic cases. Instead of changes in CCR1, our data point to a differential expression of its ligands CCL8 and CCL15 by melanoma cells as key parameters associated with metastasis and patient survival. Thus, our data suggest that melanoma-TAM interactions activates the expression of CCL8/CCL15 by melanoma cancer cells, which express CCR1, initiating an autocrine cycle of melanoma survival and invasion. Under basal conditions, CCL15 is primarily expressed in the gut and the liver and the role of CCL15-CCR1 axis has been mainly explored in hepatocellular and colorectal cancer [29]. CCL15 can induce hepatocellular carcinoma cell migration and invasion and its overexpression predicts poor patient prognosis [30, 31]. In addition to promoting tumor invasion in an autocrine manner, CCL15 also acts on non-cancer cells, which express CCR1 and CCR3, such as endothelial cells, stimulating angiogenesis; or mesenchymal stem cells, facilitating tumor niche formation [32, 33]. In immune cells, CCL15 is a potent chemoattractant for neutrophils, monocytes, and lymphocytes, signaling through CCR1 and CCR3 [34]. In cancer, CCL15 is involved in chemoattraction of CCR1 positive myeloid cells, including myeloid derived suppressor cells (MDSCs) and neutrophils, which indirectly promote primary tumor progression and metastasis in hepatocarcinoma and colorectal cancer [35-38]. Our data also point to a potential association of CCR3 positive macrophages with metastatic dissemination of primary melanomas, suggesting a paracrine role of CCL15 in promoting a particular type of prometastatic CCR3+ macrophages. To our knowledge, the role of CCL15 acting on CCR1+ melanoma cells or recruiting CCR3+ macrophages was previously unexplored in melanoma. Besides, upregulated CCL15 in melanoma may recruit CCR1+ MDSCs, as in other types of cancer [39], which are major contributors to immune checkpoint inhibitors resistance and play a crucial role in creating an immunosuppressive TME in melanoma [40]. More importantly, our findings highlight the clinical value of CCL15 content in melanoma microenvironment, which may be a useful biomarker for the prognosis of patients with primary cutaneous melanoma. The potential role of CCL15 as an independent prognostic factor warrants further prospective validation in patients that receive adjuvant therapy in order to look for predictive factors, an unmet medical need in this setting.

## Acknowledgements

We thank Alexandra de Francisco, José María Bellón and the animal facility staff for their expert technical assistance. This work was supported by the Ministry of Science and Innovation PID2021-123507OB-100 grant (PS-M, RS) co-financed by ERDF/FEDER Funds from the European Commission, “A way of making Europe”; and by Cancer Research UK, FCAECC (GCB15152947MELE) and AIRC under Accelerator Award program. CB-A and AN-V were financed by the Comunidad de Madrid YEI-program.

## Authors contribution statement

CB-A, AN-V, EG-M, AG-S and RS, performed the experiments and analyzed data; VP-B, IM-R and JAA-I, provided samples; PS-M and RS, conceived the study; RS supervised the work; CB-A, PS-M and RS wrote and edited the manuscript. All authors approved the final draft of the manuscript.

