## SUPPLEMENTAL DATA for "Chemokine profiling of melanoma-macrophage crosstalk identifies CCL8 and CCL15 as prognostic factors in cutaneous melanoma"

### SUPPLEMENTARY DATA

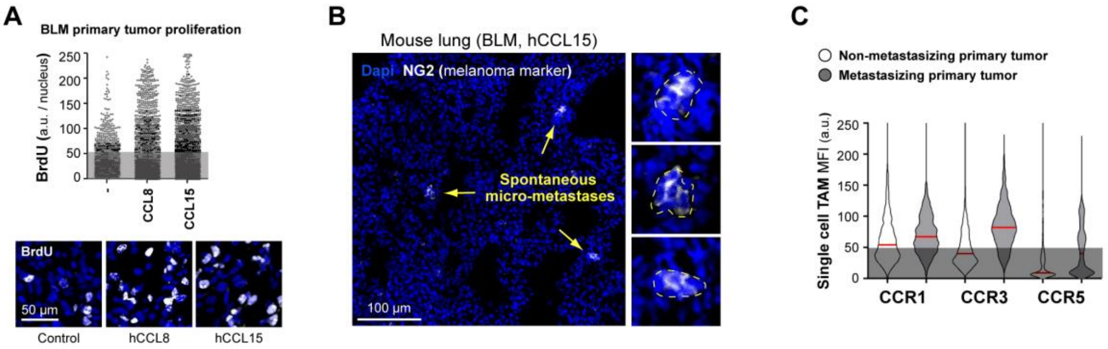


**Figure S1. Mouse model and human tissue quantifications. (A)** BrdU nuclear staining (arbitrary units, a.u.) as measured in BLM primary tumors treated with chemokines 24 h earlier (3,000 segmented nuclei are represented by condition). Representative images showing BrdU (white) within DAPI-stained nuclei (blue). **(B)** Representative images of spontaneous lung micrometastases stained with NG2 melanoma marker (white). Scale bars, 50 and 100 µm. **(C)** Single-cell MFI quantification of CCR1, CCR3 and CCR5 at segmented CD68$^{+}$ TAM (>1,300 TAMs are represented by group), as indicated.

| Gene | Primer Sequences (Roche) | Technique |
|---|---|---|
| *CCL15* | S: 5' ccaggaggatgaaggtctcc ; AS: 5' cagtggaagctttgacatcatt | qPCR |
| *CCL8* | S: 5' cccctagggacttgctcag; AS: 5' ctccagcctctggataggaa | qPCR |
| *TBP* | S: 5' cggctgtttaacttcgcttc; AS: 5' cacacgccaagaaacagtga | qPCR |

| Antigen | Antibody information | Technique |
|---|---|---|
| BrdU | R&D MAB7225 (Clone BU-1), mouse IgG2A | Immunoflourescence (IF) |
| NG2 | BD Pharmingen 554275 (Clone 9.2.27), mouse IgG2A | Immunoflourescence (IF) |
| CCL15 | R&D MAB363 (Clone 52130), mouse IgG1 | IF-paraffin (pH 9.0) |
| CCL8 | Biorbyt orb13564, Polyclonal rabbit | IF-paraffin (pH 9.0) |
| CCR1 | R&D MAB145 (Clone 53504), mouse IgG2b | IF-paraffin (pH 9.0); Invasion blockade |
| CCR1 | Thermofisher PA1-41062, Polyclonal rabbit | Western Blotting |
| CCR3 | R&D MAB155 (Clone 61828), rat IgG2A | IF-paraffin (pH 9.0); Invasion blockade |
| CCR3 | abcam (Clone Y31), rabbit | Western Blotting |
| CCR5 | R&D MAB182 (Clone 45531), mouse IgG2b | IF-paraffin (pH 9.0); Invasion blockade |
| CCR5 | Santa Cruz sc-17833 (Clone D-6), mouse IgG1 | Western Blotting |
| CD68 | Dako M087601-2 (Clone PG-M1), mouse IgG3 | IF-paraffin (pH 9.0) |
| Isotype Control | Biolegend #401202, mouse IgG2b | Invasion blockade |
| Isotype Control | eBioscience #14-4321-82, rat IgG2a | Invasion blockade |

**Table S1. Primers and antibodies.**

| | Ratio BLM+M1 to M1 (Log2) | Ratio BLM+M1 to BLM (Log2) | Ratio BLM+M2 to M2 (Log2) | Ratio BLM+M2 to BLM (Log2) |
|---|---|---|---|---|
| **CCL15** | 86.7 | 429.5 | 297 | 317.5 |
| **CCL5** | 9.6 | 3.3 | 71.6 | 11.6 |
| **CXCL1** | 6.3 | 0.9 | 61.4 | 6.5 |
| **CCL20** | 4.8 | 21.7 | 50.4 | 4.2 |
| **CCL8** | 4 | 20.4 | 30 | 4.3 |
| **CCL13** | 2.8 | 12.7 | 16.7 | 3.4 |
| **CCL1** | 2.7 | 35.9 | 15.9 | 7 |
| **CCL7** | 2.4 | 10.1 | 15.8 | 19.7 |
| **CXCL6** | 1.9 | 6.5 | 13.9 | 3.7 |
| **CXCL10** | 1.9 | 5.6 | 11.5 | 4.3 |
| **CXCL5** | 1.6 | 1.5 | 9.9 | 3.6 |
| **CCL3** | 1.5 | 29.8 | 9.9 | 3.9 |
| **CCL4** | 1.4 | 5.3 | 9.5 | 3.7 |
| **CXCL7** | 1.3 | 6.3 | 6.5 | 0.8 |
| **CCL23** | 1.1 | 28.4 | 6.2 | 6.5 |
| **CCL2** | 1.1 | 1.2 | 6.1 | 2.5 |
| **CCL16** | 1 | 3.8 | 5.9 | 1.4 |
| **CCL24** | 1 | 2.2 | 5.6 | 5.6 |
| **CCL17** | 1 | 4.5 | 4.5 | 15.6 |
| **CXCL4** | 1 | 3.6 | 4.3 | 10.3 |
| **CCL18** | 1 | 3.4 | 3.9 | 0.8 |
| **CXCL8** | 0.9 | 0.9 | 3.7 | 5.5 |
| **CXCL16** | 0.9 | 4.2 | 3.5 | 33.8 |
| **CXCL12** | 0.9 | 2.8 | 3.1 | 3.7 |
| **XCL1** | 0.9 | 3.5 | 2.4 | 3.6 |
| **CCL22** | 0.9 | 1.7 | 2.3 | 3.6 |
| **CXCL11** | 0.8 | 3.3 | 2.1 | 2.7 |
| **CCL26** | 0.7 | 3.8 | 2 | 1.8 |
| **CCL14** | 0.7 | 3.8 | 1.7 | 1.8 |
| **CCL19** | 0.7 | 4 | 1.5 | 4.4 |
| **CCL27** | 0.6 | 3 | 1.3 | 3.8 |
| **CCL28** | 0.6 | 4.3 | 1.1 | 0.9 |
| **CCL21** | 0.5 | 2.8 | 0.9 | 1.1 |
| **CCL25** | 0.1 | 5.8 | 0 | 30 |
| **CCL11** | 0 | 0 | 0 | 4.7 |
| **CXCL9** | 0 | 0 | 0 | 0 |
| **CXCL12** | 0 | 1.2 | 0 | 0 |
| **CXCL13** | 0 | 0 | 0 | 0 |

**Table S2. Secreted chemokine expression.** List of the 38 secreted chemokines quantified in 72 hours supernatants of macrophages + BLM melanoma cells *vs* macrophages or BLM cells alone (n= 4 donors).

| | | TAM CCL15 | P | TC CCL15 | P | TAM CCL8 | P | TC CCL8 | P | TAM CCR3 | P |
|---|---|---|---|---|---|---|---|---|---|---|---|
| **Gender** | Male (n=26) | 75±27 | 0.8584 | 98±26 | 0.727 | 107±31 | 0.1356 | 88±33 | 0.1756 | 110±43 | 0.0095 |
| | Female (n=36) | 77±26 | | 102±42 | | 92±27 | | 76±29 | | 79±32 | |
| **Age** (years) | ≤65 (n=27) | 79±30 | 0.4392 | 94±32 | 0.2722 | 102±31 | 0.5267 | 89±27 | 0.0897 | 92±40 | 0.954 |
| | >65 (n=40) | 72±24 | | 104±38 | | 94±31 | | 73±34 | | 91±39 | |
| **Location** | Head/limbs (n=41) | 69±21 | 0.0333 | 94±35 | 0.0777 | 91±36 | 0.4542 | 78±35 | 0.5763 | 89±38 | 0.5406 |
| | Trunk (n=26) | 85±31 | | 107±35 | | 102±26 | | 82±28 | | 95±41 | |
| **Subtype** | Nodular (n=27) | 77±28 | 0.9287 | 107±41 | 0.2808 | 97±29 | 0.9874 | 78±26 | 0.6842 | 101±39 | 0.1691 |
| | Others (n=40) | 74±25 | | 94±31 | | 97±34 | | 81±35 | | 86±39 | |
| **Ulceration** | No (n=34) | 75±26 | 0.6993 | 94±37 | 0.3678 | 90±29 | 0.1234 | 82±28 | 1 | 85±36 | 0.0975 |
| | Yes (n=29) | 73±24 | | 102±31 | | 103±34 | | 79±35 | | 102±41 | |
| **Breslow** | ≤4mm (n=34) | 74±30 | 0.4096 | 101±41 | 0.9107 | 91±28 | 0.1831 | 76±33 | 0.2977 | 87±36 | 0.6626 |
| | >4mm (n=33) | 76.5±24 | | 98±31 | | 104±34 | | 83±31 | | 94±41 | |
| **Stage** | II (n=48) | 75±28 | 0.6511 | 95±32 | 0.1693 | 97±28 | 0.9156 | 79±29 | 0.5497 | 89±36 | 0.7063 |
| | III-IV (n=19) | 76±22 | | 111±42 | | 96±41 | | 82±38 | | 98±46 | |
| **Metastasis** | Non met (n=31) | 69±24 | 0.0273 | 81±21 | P<0.0001 | 88±31 | 0.101 | 71±32 | 0.0243 | 77±39 | 0.0062 |
| | Met (n=36) | 81±28 | | 116±38 | | 105±30 | | 87±29 | | 104±36 | |
| **Survival** | Yes (n=44) | 69±24 | 0.0121 | 89±28 | 0.0004 | 97±33 | 0.9437 | 73±33 | 0.0043 | 87±42 | 0.1469 |
| | No (n=23) | 86±27 | | 121±41 | | 98±28 | | 96±20 | | 99±34 | |

**Table S3. Association of evaluated prognostic markers with clinicopathological features (Mann-Whitney).**
